# The role of ion dissolution in metal and metal oxide surface inactivation of SARS-CoV-2

**DOI:** 10.1101/2023.09.08.556901

**Authors:** Jane Hilton, Yoshiko Nanao, Machiel Flokstra, Meisam Askari, Terry K. Smith, Andrea Di Falco, Phil D.C. King, Peter Wahl, Catherine S Adamson

## Abstract

Antiviral surface coatings are under development to prevent viral fomite transmission from high-traffic touch surfaces in public spaces. Copper’s antiviral properties have been widely documented; but the antiviral mechanism of copper surfaces is not fully understood. We screened a series of metal and metal oxide surfaces for antiviral activity against severe acute respiratory syndrome coronavirus 2 (SARS-CoV-2), the causative agent of coronavirus disease (COVID-19). Copper and copper oxide surfaces exhibited superior anti-SARS-CoV-2 activity; however, level of antiviral activity was dependent upon the composition of the carrier solution used to deliver virus inoculum. We demonstrate that copper ions released into solution from test surfaces can mediate virus inactivation, indicating a copper ion dissolution-dependent antiviral mechanism. Level of antiviral activity is, however, not dependent on the amount of copper ions released into solution *per se*. Instead, our findings suggest that degree of virus inactivation is dependent upon copper ion complexation with other biomolecules (e.g., proteins/metabolites) in the virus carrier solution that compete with viral components. Although using tissue culture-derived virus inoculum is experimentally convenient to evaluate the antiviral activity of copper-derived test surfaces, we propose that the high organic content of tissue culture medium reduces the availability of “uncomplexed” copper ions to interact with the virus, negatively affecting virus inactivation and hence surface antiviral performance. We propose that laboratory antiviral surface testing should include virus delivered in a physiologically relevant carrier solution (saliva or nasal secretions when testing respiratory viruses) to accurately predict real-life surface antiviral performance when deployed in public spaces.

**Importance:** The purpose of evaluating antiviral activity of test surfaces in the laboratory is to identify surfaces that will perform efficiently in preventing fomite transmission when deployed on high-traffic touch surfaces in public spaces. The conventional method in laboratory testing is to use tissue culture-derived virus inoculum, however this study demonstrates that antiviral performance of test copper-containing surfaces is dependent on the composition of the carrier solution in which the virus inoculum is delivered to test surfaces. Therefore, we recommend that laboratory surface testing should include virus delivered in a physiologically relevant carrier solution, to accurately predict real-life test surface performance in public spaces. Understanding the mechanism of virus inactivation is key to future rational design of improved antiviral surfaces. Here, we demonstrate that copper ions released from copper surfaces into small liquid droplets containing SARS-CoV-2, is a mechanism by which the virus that causes COVID-19 can be inactivated.

## Introduction

Antiviral surface coatings are a non-pharmacological intervention that aim to prevent virus transmission via virus-contaminated surfaces, termed fomites (1). Fomite transmission occurs by hand contamination through touching fomites and subsequent self-inoculation by transfer of infectious virus from contaminated hands to exposed mucosal membranes in the mouth, nose, and eyes. Fomites are typically high-traffic touch surfaces, such as handles, push plates, lift buttons, railings, telephones, touch screens, counter tops etc., located in a wide range of public spaces. Particularly notable are ones located in healthcare settings such as hospitals and care homes. Fomite transmission plays an important role in the spread of enteric and respiratory viruses (2), although both groups of viruses have more than one route of transmission. Like other respiratory viruses, SARS-CoV-2, the causative agent of the COVID-19 pandemic, is primarily transmitted via droplet/aerosol mediated airborne transmission, but fomites are also considered a mode of SARS-CoV-2 transmission by the World Health Organisation (https://www.who.int/news-room/questions-and-answers/item/coronavirus-disease-covid-19-how-is-it-transmitted) and others (3, 4).

Fomite transmission requires that the virus remains viable on a surface long enough for onward human transfer. Laboratory studies have shown that SARS-CoV-2 remains viable on a timescale ranging from hours to days on a variety of commonly used surface materials such as stainless steel, plastic, and paper (5, 6). Environmental studies have shown that SARS-CoV-2 RNA has been detected on a wide range of surfaces in public spaces, particularly medical settings (4), however studies attempting to detect viable virus from such environmental surface samples are scarce and technically demanding. As a result, these studies often fail to detect viable virus (7, 8), however evidence that viable SARS-CoV-2 can be recovered from fomites in real-life settings has been reported (4, 9, 10). Longevity of viral surface survival is an important parameter affecting the likelihood of fomite transmission; prevention of fomite transmission depends on rapid inactivation of viruses on surfaces that act as fomites.

Surface survival times are dependent on the virus, environmental factors (e.g., temperature, humidity) and surface properties (e.g., chemical composition, porosity) (2, 4). Development of surfaces with antiviral properties offers a long-term behaviour-independent strategy to prevent fomite transmission, as opposed to commonly employed short-term behaviour-dependent strategies including frequent hand washing and surface disinfection regimens. Copper and its alloys have long been known for their antimicrobial properties and laboratory studies of copper surfaces have been shown to inactivate a wide range of viruses, bacteria and fungi (11–13). The mechanism by which copper surfaces inactivate pathogens has not been fully elucidated, however with respect to viruses two key mechanisms have been proposed; (i) direct contact between the virus and the solid copper-containing surface (copper dissolution independent) and/or (ii) ion dissolution resulting in release of copper ions into solution from the copper-containing surface (copper dissolution dependent) (12). Virus inactivation has been reported to occur via damage to viral proteins, genomic material and envelopes (12).

In this study, we screened a series of metal and metal oxide surfaces for antiviral activity against SARS-CoV-2. Copper and copper oxide surfaces exhibited superior anti-SARS-CoV-2 activity; however, the level of antiviral activity was dependent upon composition of the carrier solution used to deliver virus inoculum. We demonstrate that copper ion dissolution is a mechanism of SARS-CoV-2 inactivation, but it is not dependent upon the amount of total copper ions released into solution per se. Instead, we suggest that the degree of virus inactivation is dependent upon copper ion complexation with other biomolecules in the virus inoculum that compete with viral components. Based on our findings, we recommend that laboratory antiviral surface testing should include virus delivered in a physiologically relevant carrier solution (i.e., saliva or nasal secretions when testing respiratory viruses) to predict real-life test surface performance more accurately when deployed in public spaces.

## Results

### SARS-CoV-2 is inactivated upon exposure to copper surfaces

The antiviral properties of copper are confirmed against SARS-CoV-2, by its time dependent inactivation upon exposure to bulk copper foil or thin-film evaporated copper test coupons (Fig. 1). Significantly less virus inactivation occurred upon exposure to stainless steel and the virus remains consistently viable across the time series with respect to the no coupon control. SARS-CoV-2 inactivation was essentially comparable between the two types of copper samples studied. No viable virus was detectable after 120-min exposure to either copper surface, demonstrating that exposure to a copper surface requires at least 1-2 hours to efficiently inactivate the virus inoculum used (∼ 4,000 PFU/7 μL in Dulbecco’s Modified Eagle’s Medium supplemented in 2% v/v FBS (DMEM-2%FBS)). We applied an exponential fit to our data to determine the mean half-life of the virus when exposed to copper surfaces (Fig. 1B), which was 38 and 28 minutes for the copper foil and evaporated copper surfaces respectively. Therefore, a 30-min exposure to a copper surface results in ∼50% virus inactivation (Fig. 1A), providing an ideal time point to screen further test coupons to identify surface materials that inactivate SARS-CoV-2 faster than copper and thus demonstrate improved antiviral properties.

**FIG 1.**
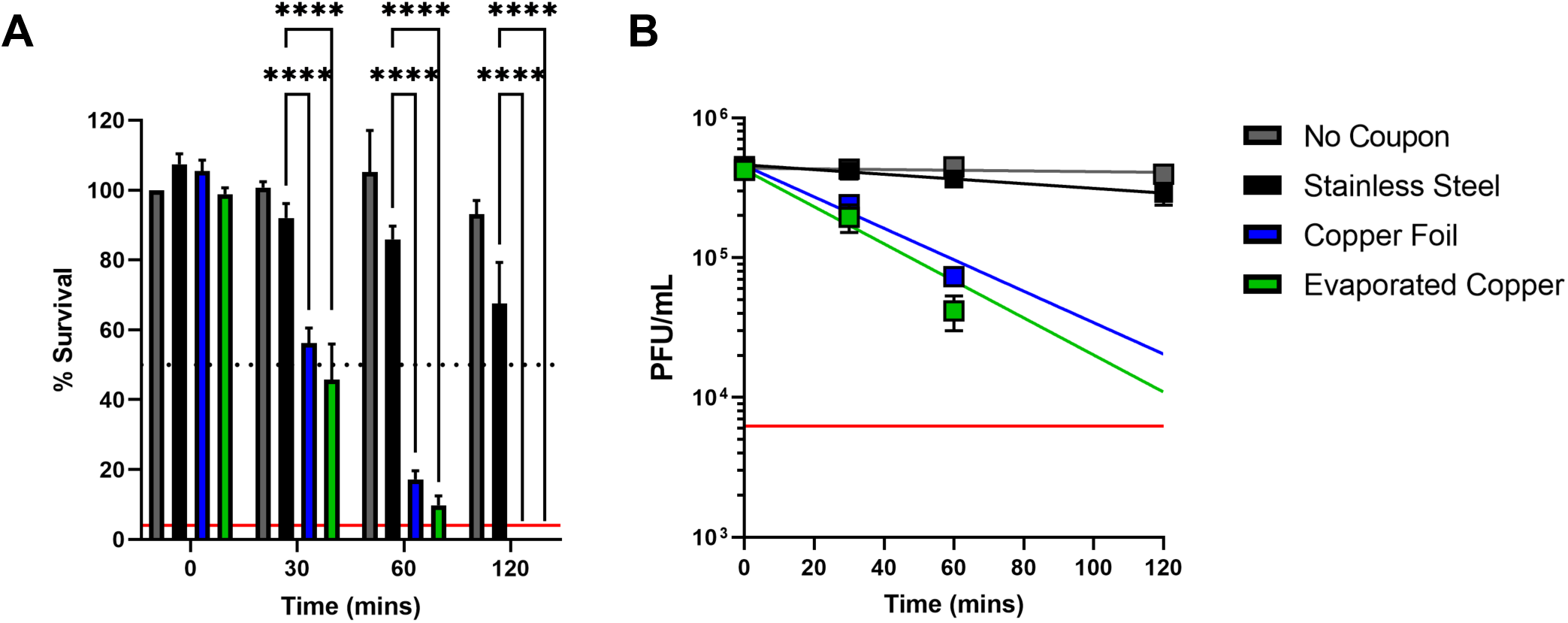
SARS-CoV-2 inactivation upon exposure to copper surfaces over time. (A) Percent survival of SARS-CoV-2 exposed to different metal surfaces after 0-, 30-, 60-and 120-min. Data is expressed as a percentage of a no coupon control at 0-min time point for each test condition. Data shown represents mean values (n = 3 replicates and error bar = SD) and is representative of 3 independent experiments. Statistical significance was assessed using two-way ANOVA with Tukeys multiple comparison test, **** p < 0.0001. The limit of detection (LOD) for the assay is indicated by the solid red line and 50% inactivation is indicated by the black dotted line. (B) Titre of SARS-CoV-2 (PFU/mL) exposed to different test surfaces as a function of time, exponential fits to the data are shown along with a solid red line, which indicates the LOD for the assay.

### Screening metal and metal oxide surfaces revealed that Cu_2_O containing surfaces exhibited SARS-CoV-2 antiviral activity superior to elemental copper

Utilizing the 30-min copper exposure time point as a screening reference point, a selection of different surfaces were tested with the aim of identifying materials that exhibit antiviral activity superior to copper. The thin-film evaporated copper coupon (500 nm thickness) was chosen as the standard reference point (referred to as copper), along with stainless steel and no coupon controls, for screening purposes and throughout the manuscript. Screening was performed using the same SARS-CoV-2 inoculum described above (∼ 4,000 PFU/7 μL in DMEM-2%FBS).

Initially, we generated a series of coupons with elemental metal surfaces; transition metals silver (Ag), nickel (Ni) and palladium (Pd) were selected based on their proximity to copper in the periodic table, along with post-transition metal bismuth (Bi) (Table S1 and S2). Upon exposure of SARS-CoV-2 to the test elemental metal surfaces it was clearly apparent that copper exhibited the best antiviral activity (Fig. 2A). We next investigated the antiviral properties of transition metal oxide surfaces. In the first instance, we generated delafossite copper chromate (CuCrO_2_), titanium oxide (TiO_2_) and indium tin oxide (ITO) films (Table S1 and S2). ITO was particularly selected as a transparent conductor, with widespread application in touch screen surfaces. Unfortunately, these test surfaces did not result in any substantial SARS-CoV-2 inactivation and again copper exhibited the best antiviral activity (Fig. 2B).

**FIG 2.**
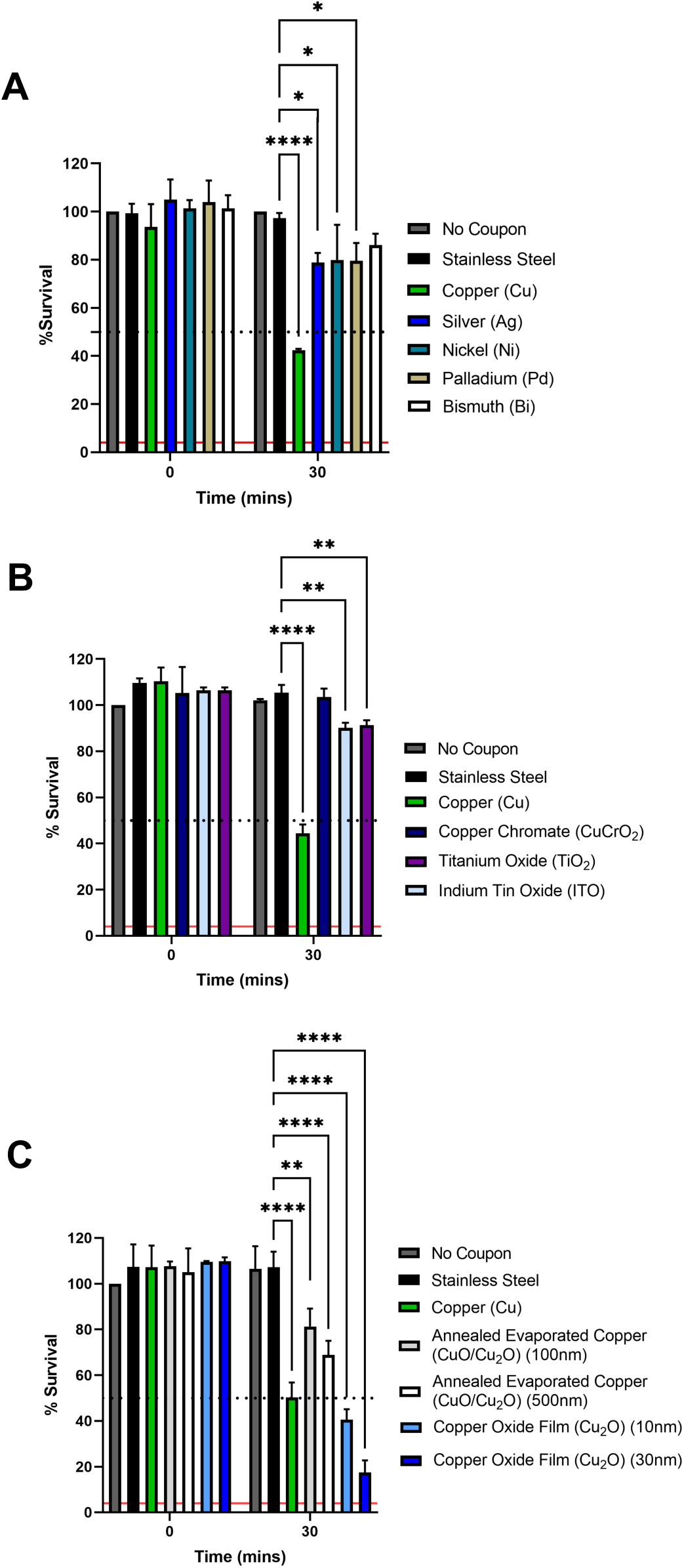
Screening test elemental metal and metal oxide surfaces for SARS-CoV-2 antiviral activity superior to copper. Percent survival of SARS-CoV-2 exposed to different test metal and metal oxide surfaces after 0 and 30 min compared to no coupon, stainless steel, and copper controls. Data is expressed as a percentage of a no coupon control at 0 min time point for each test condition. (A) elemental metal test surfaces; silver (Ag), nickel (Ni), palladium (Pd), bismuth (Bi) (B) metal oxide test surfaces; copper chromate (CuCrO_2_), titanium oxide (TiO_2_), indium tin oxide (ITO) and (C) copper oxide test surfaces; annealed evaporated copper (CuO/Cu_2_O mixture) and predominantly Cu_2_O containing surfaces, generated at indicated thicknesses. Data shown represents mean values (n = 3 replicates and error bar = SD) and is representative of 3 independent experiments. Statistical significance was assessed using two-way ANOVA with Tukeys multiple comparison test, **** p < 0.0001, ** p < 0.01, * p < 0.1. The limit of detection (LOD) for the assay is indicated by the solid red line and 50% inactivation is indicated by the black dotted line.

Given that copper consistently exhibited the best antiviral activity, we proceeded to test copper oxide surfaces. Using two different methods, we generated two types of copper oxide surfaces: (i) an annealed mixture of cupric oxide and cuprous oxide (CuO/Cu_2_O), and (ii) a copper oxide thin film consisting predominantly of cuprous oxide (Cu_2_O) (Table S1 and Fig.S1). Each type of copper oxide surface was generated at two different thicknesses (Table S1). The copper oxide surfaces all exhibited significant antiviral activity (Fig. 2C). Importantly, the Cu_2_O thin-film exhibited better virus inactivation than annealed copper surfaces containing a CuO/Cu_2_O mixture in the surface layer, suggesting that the Cu_2_O oxidation state has superior antiviral properties. Most notably, the Cu_2_O thin-films resulted in better virus inactivation than the copper reference coupon, with the thicker (∼30 nm) Cu_2_O film resulting in ∼75% SARS-CoV-2 inactivation after 30 min exposure, which represents an improvement of inactivation by ∼50% compared to copper. For the mixed CuO/Cu_2_O samples, we found that the more oxygen-rich CuO phase forms as the surface layer, with Cu_2_O forming below the surface (Fig. S2), inhibiting the superior antiviral properties of Cu_2_O. Interestingly, we also observed increased antiviral activity for the thicker (∼30 nm) copper oxide films, with a somewhat reduced inactivation for the ultrathin (∼10 nm) film thickness. Overall, we show that thin films exposing Cu_2_O at the surface have superior antiviral properties over an evaporated and post-oxidized copper surface and that for films with a thickness of tens of nanometers, the film thickness can limit the degree of antiviral activity observed.

### Increasing copper surface thickness correlates with increased SARS-CoV-2 inactivation

Motivated by these findings of a thickness-dependent antiviral activity of copper oxide films, we took advantage of our ability to precisely control film thickness by generating a series of evaporated copper films with thicknesses of 5, 10, 20, 50, 100, 250 and 500 nm. In agreement with our previous observations, the amount of SARS-CoV-2 inactivation after 30-min exposure increased stepwise with film thickness from 5-50 nm and stabilized at ∼50% inactivation when exposed to copper films of ≥50 nm (Fig. 3A). The stabilization at ≥50 nm is likely to be a function of the 30-min copper exposure time, as we demonstrated in Fig. 1A, where further inactivation occurs after 60- and 120-min exposure to a 500 nm copper film. We also observed that following removal of 7 μL in DMEM-2%FBS after 30-min exposure time, the copper film appeared modified on the coupons generated with a copper film thickness of 50 nm, whereas the copper film remained visible on coupons with a 500 nm layer (Fig. 3B). These observations, combined with the fact that increasing copper film thickness correlates with increased SARS-CoV-2 inactivation, suggests that dissolution of copper ions into solution might be the mechanism driving virus inactivation.

**FIG 3.**
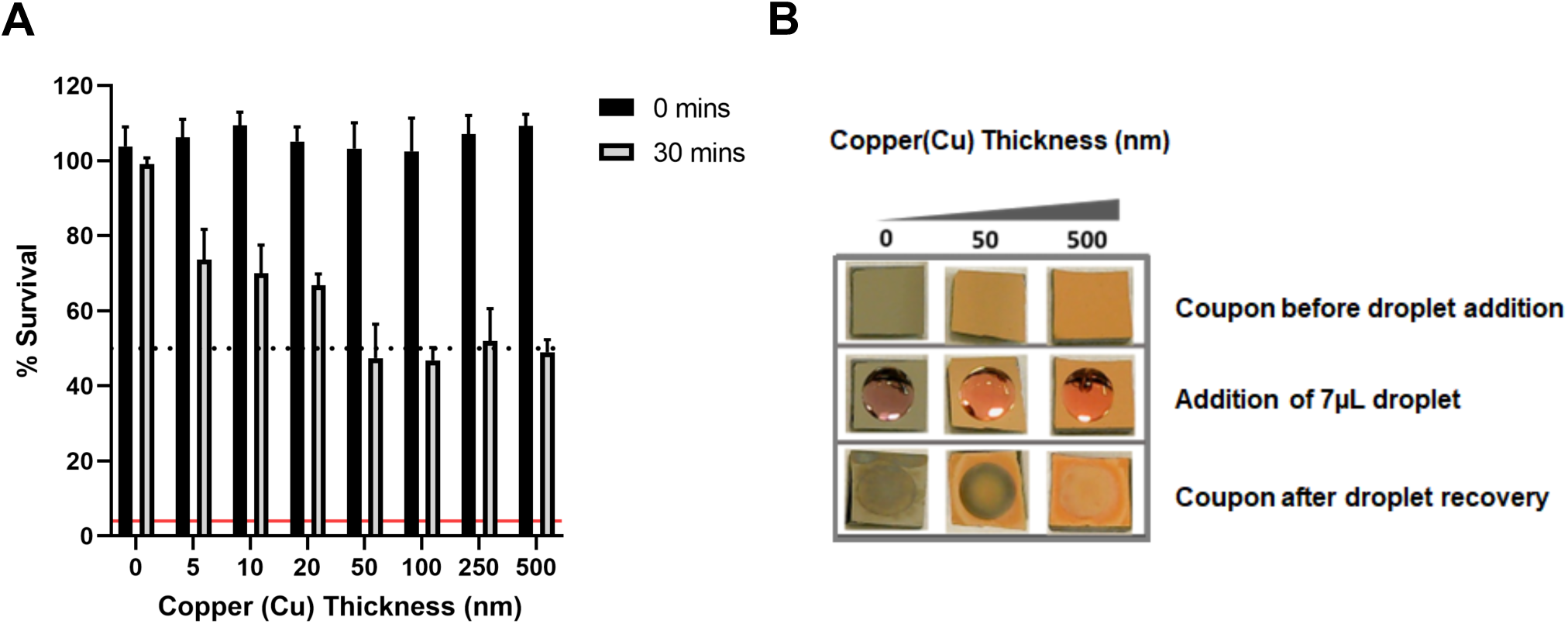
Effect of copper surface film thickness on SARS-CoV-2 inactivation. (A) Percent survival of SARS-CoV-2 exposed to coupons with evaporated copper film of increasing thickness. Data shown represents mean values (n = 3 replicates and error bar = SD) and is representative of 3 independent experiments. The limit of detection (LOD) for the assay is indicated by the solid red line and 50% inactivation is indicated by the black dotted line. (B) images of evaporated copper thin-film coupons of 50 nm and 500 nm thicknesses before, during and after 30 min incubation with a 7 μL droplet of DMEM-2%FBS.

### Different carrier solutions impact SARS-CoV-2 inactivation, but inactivation does not correlate with amount of Cu ions released into solution

Understanding the mechanism of virus inactivation is key to future rational design of improved antiviral surfaces. Although the antiviral mechanism remains poorly understood two main hypotheses have been proposed; (i) direct contact between the virus and the solid copper-containing surface (copper dissolution independent) and/or (ii) ion dissolution resulting in release of copper ions into solution from the copper-containing surface (copper dissolution dependent) (12). To further investigate the role of copper ion dissolution we hypothesized that if the virus was delivered to a test copper surface in carrier solutions that differentially dissolve copper ions, then virus inactivation would be correspondingly affected. We selected the following carrier solutions; DMEM-2%FBS, phosphate buffered saline (PBS) and artificial saliva (AS). DMEM-2%FBS is equivalent to the virus inoculum used in our prior experiments, PBS is a physiological buffered solution commonly used in cell culture and AS was selected to simulate a real-world scenario related to transmission of respiratory viruses such as SARS-CoV-2.

ICP-OES (Inductively Coupled Plasma - Optical Emission Spectroscopy) was used to determine Cu ion concentration released into 7 μL of each carrier solution following a 30-min exposure to either reference evaporated copper (500 nm) or Cu_2_O (30 nm) containing thin-film coupons (Fig. 4A). The largest amount of Cu ion dissolution was observed upon DMEM-2%FBS exposure to evaporated copper followed by Cu_2_O containing coupons. Approximately one third the level of copper ions was released upon PBS exposure for both coupon types and the least amount was observed upon AS exposure, which resulted in a low-level ion release from the Cu_2_O coupon and no detectable release of copper ions for the evaporated copper coupon.

**FIG 4.**
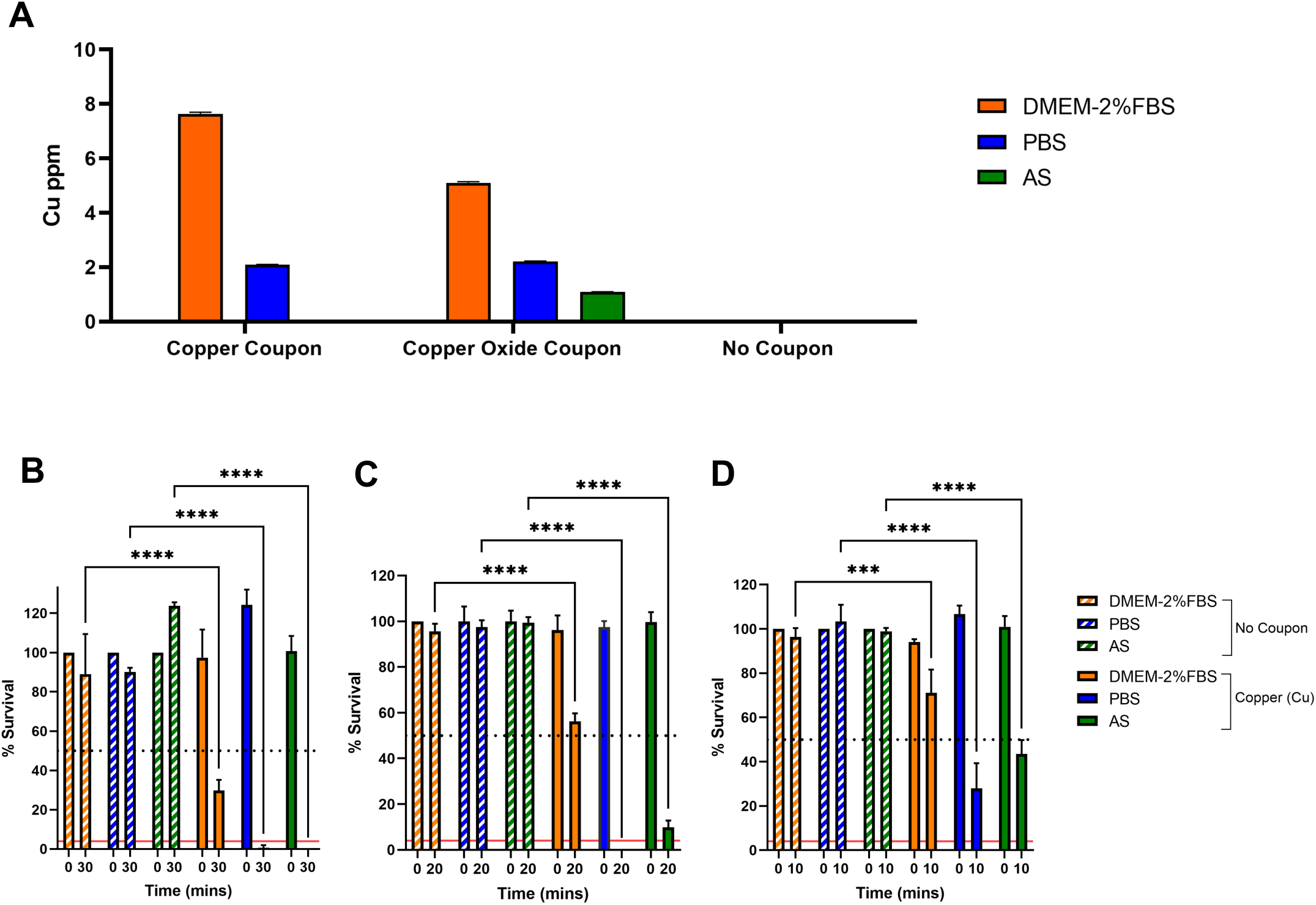
Impact of different carrier solutions on copper ion dissolution and SARS-CoV-2 inactivation upon exposure to an evaporated copper thin-film surface. (A) ICP-OES determined copper ion levels in DMEM-2%FBS, PBS or AS carrier solutions following 30-mi exposure to evaporated copper, Cu_2_O thin film coupons or no coupon control. Data shown represents mean values (n = 6 replicates and error bar = SD). (B-D) percent survival of SARS-CoV-2 resuspended in DMEM-2%FBS, PBS or AS carrier solutions and exposed to evaporated copper surfaces for (B) 30, (C) 20 and (D) 10 min or the equivalent no coupon control. Data is expressed as a percentage of a no coupon control at 0 min time point for each test condition. Data shown represents mean values (n = 3 replicates and error bar = SD). At the 30 min time point the data shown is representative of 3 independent experiments, the 20- and 10-min time points were included in the 3^rd^ and final experimental repeat. Statistical significance was assessed using two-way ANOVA with Tukeys multiple comparison test, **** p < 0.0001, *** p < 0.001. The limit of detection (LOD) for the assay is indicated by the solid red line and 50% inactivation is indicated by the black dotted line.

We next tested virus inactivation following exposure to evaporated copper coupons when SARS-CoV-2 is delivered as an inoculum of ∼4,000 PFU in 7 μL of each carrier solution. Importantly, we confirmed that SARS-CoV-2 remained comparably viable in each carrier solution; this was tested by measuring SARS-CoV-2 titre after resuspension in each carrier solution to confirm equal virus input (Fig. S3) and is demonstrated by the virus remaining consistently viable across the time series for each carrier solution with respect to the no coupon control (Fig. 4B-D). At the previously used 30-min exposure time, SARS-CoV-2 in DMEM-2%FBS resulted in ∼70% inactivation, unexpectedly however viable virus was undetectable when SARS-CoV-2 was delivered in either PBS or AS (Fig. 4B). On the 3^rd^ and final repeat of this experiment we conducted coupon exposure at reduced time points of 20- and 10-mins, reassuringly the level of SARS-CoV-2 inactivation was time dependent for each carrier solution (Fig. 4C and D). Overall, we show that virus inactivation is impacted by virus carrier solution and the most effective inactivation occurred when virus was delivered in PBS. However, the level of SARS-CoV-2 inactivation does not appear to correlate with the amount of available copper ions in the presence of the different virus carrier solutions.

### Copper ion dissolution and availability is a mechanism that can independently lead to SARS-CoV-2 inactivation

To directly test the role of copper ion dissolution in SARS-CoV-2 inactivation, we performed a variation of the test surface inactivation assay, in which virus inactivation is de-coupled from the test copper surface. First, 7 μL of each carrier solution was added to either evaporated copper or Cu_2_O containing thin-film coupons and incubated for 0- or 30-mins. The carrier solution (together with any released copper ions) was then removed from the test surface and spiked with 2 μL SARS-CoV-2 inoculum containing ∼4,000 PFU and further incubated for 0- or 30-mins. To act as a control, we performed a test surface inactivation assay (Fig. 5A and B) in parallel to the de-coupled assay (Fig. 5C and D).

**FIG 5.**
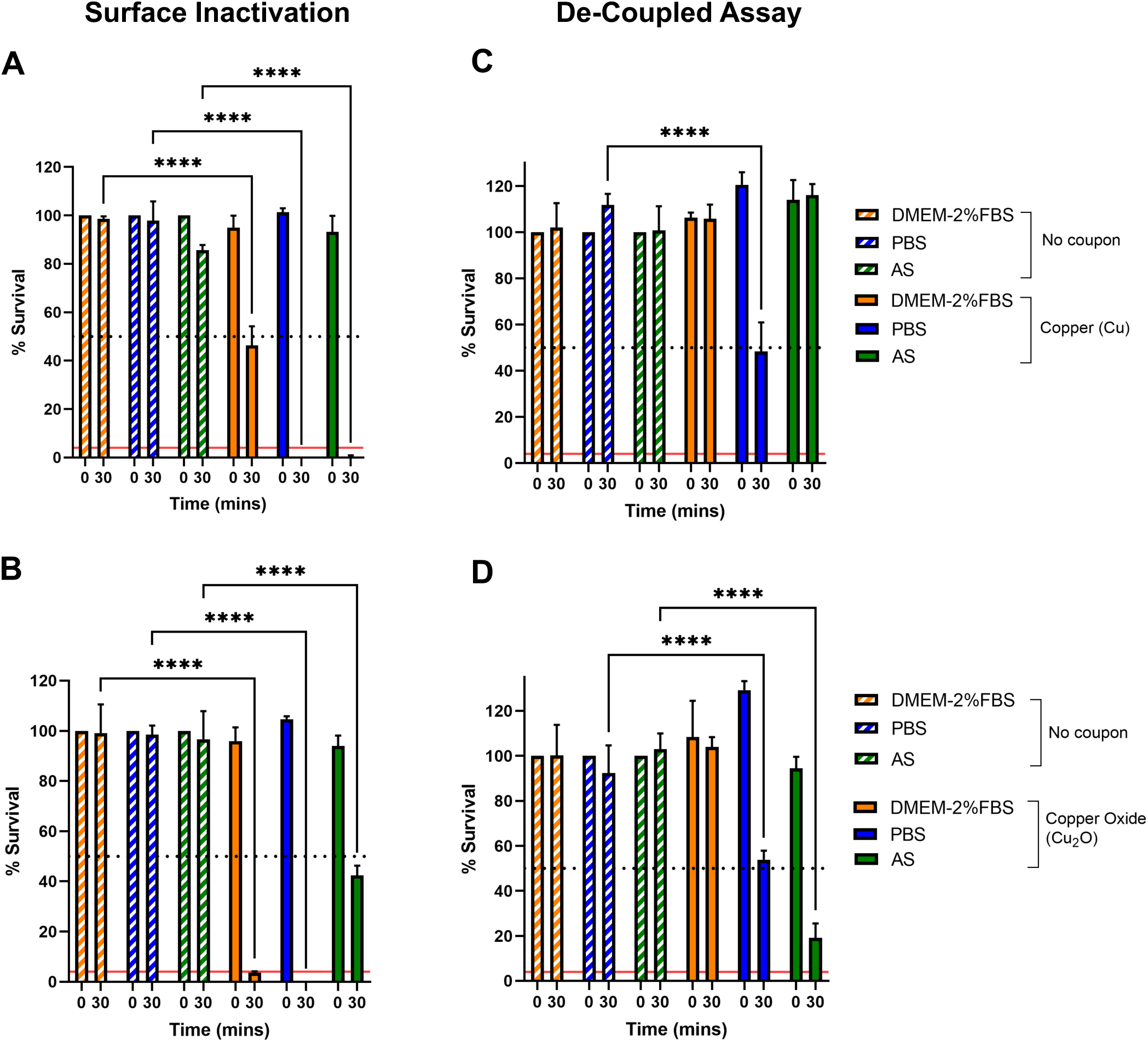
De-coupled ion dissolution SARS-CoV-2 inactivation assay. (A and B) test surface virus inactivation assay: percent survival of SARS-CoV-2 resuspended in DMEM-2%FBS, PBS or AS carrier solutions and exposed to (A) evaporated copper or (B) Cu_2_O thin-film coupons for 0 or 30 minutes or the equivalent no coupon control. (C and D) de-coupled virus inactivation assay: carrier solution DMEM-2%FBS, PBS or AS exposed to evaporated copper (C) and Cu_2_O (D) thin-film coupons for 0 or 30 min or the equivalent no coupon control. Following coupon exposure, the resultant solution is removed and spiked with SARS-CoV-2 and incubated for a further 0 or 30 min or the equivalent no coupon control. Data is shown as percent survival of SARS-CoV-2 is expressed as a percentage of a no coupon control at 0-min time point for each test condition. Data represents mean values (n = 3 replicates and error bar = SD) and is representative of 3 independent experiments. Statistical significance was assessed using two-way ANOVA with Tukeys multiple comparison test, **** p < 0.0001. The limit of detection (LOD) for the assay is indicated by the solid red line and 50% inactivation is indicated by the black dotted line.

As expected from the result shown in Fig. 2C, 30-min exposure of SARS-CoV-2 in DMEM-2%FBS to either an evaporated copper or Cu_2_O coupon resulted in ∼50% and ∼90% inactivation respectively (Fig. 5A and B). In agreement with the results described in Fig. 4B, 30-min exposure of SARS-CoV-2 in PBS or AS to an evaporated copper coupon resulted in 100% inactivation (Fig. 5A). However, upon exposure to a Cu_2_O surface SARS-CoV-2 in AS only resulted in 50% inactivation (Fig. 5B), thus the presence of the Cu_2_O did not result in the improved virus inactivation observed when virus is delivered in DMEM-2%FBS. In fact, superior inactivation occurred when SARS-CoV-2 is delivered in AS and exposed to the evaporated copper surface (Fig. 5A and B).

The de-coupled assay, which directly tests if copper ions released into solution can inactivate virus, showed that the DMEM-2%FBS-based solution recovered from either evaporated copper or Cu_2_O containing surfaces was not capable of any SARS-CoV-2 inactivation (Fig. 5C and D), despite the ICP-OES analysis demonstrating that the greatest level of copper ions is released when coupons are exposed to DMEM-2%FBS (Fig. 4A). In contrast, the PBS-based solution recovered from either surface was capable of ∼50% virus inactivation (Fig. 5C and D) providing evidence that copper ion dissolution, and hence Cu ion released into solution, can be a mechanism directly and independently responsible for virus inactivation, but no advantage was afforded by release of ions from the Cu_2_O film. Curiously, the AS-based solution recovered from the evaporated copper surface was not capable of virus inactivation (Fig. 5C), yet the AS-based solution recovered from the Cu_2_O containing surface resulted in the most potent virus inactivation (∼80%) observed for the de-coupled assay and surprisingly was even better than the level of inactivation when virus in AS was in direct contact with the Cu_2_O containing surface (Fig. 5D). Overall, we show that copper ions resulting from dissolution independent of the direct surface contact is a mechanism that can independently lead to SARS-CoV-2 inactivation, but this mechanism is influenced by the properties of the carrier solution and the type of copper ions.

## Discussion

Copper has been widely documented to exert antiviral activity however, the mechanism of action is not fully understood. In this study we further investigate the role of ion dissolution as a mechanism by which copper and copper oxide surfaces inactivate SARS-CoV-2. First, we confirmed that SARS-CoV-2 is efficiently inactivated upon exposure to copper surfaces, in broad agreement with other SARS-CoV-2 studies (5, 14–18).

We screened a series of metal coupons with the aim of identifying a surface that inactivates SARS-CoV-2 faster than copper. Despite antimicrobial properties of silver being widely reported (19), we show that a silver surface did not exhibit extensive SARS-CoV-2 inactivation. Others have also reported silver materials to lack antiviral activity against SARS-CoV-2 and other viruses (14, 16, 20, 21) and the poor antiviral activity has been proposed to be due to low levels of Ag ion dissolution (14, 20). In contrast, positive reports of silver antiviral activity generally relate to silver-containing nanoparticles (22–26). The other elemental metals tested in this study (nickel, palladium and bismuth) also exhibited weak antiviral activity against SARS-CoV-2.

We tested a series of transition metal oxide surfaces. A titanium oxide (TiO_2_) surface did not result in a substantial level of SARS-CoV-2 inactivation. TiO_2_ has photocatalytic properties that following light illumination generates highly oxidizing free radicals (reactive oxygen species) that are reported to have antibacterial and antiviral activity (27). Our experimental procedure did not include a deliberate illumination step; however, it has been reported that when illumination of TiO_2_ or TiO_2_-composite surface coatings is undertaken, significant levels of SARS-CoV-2 inactivation occur (28–32). Weak anti-SARS-CoV-2 activity was observed upon exposure to an indium tin oxide (ITO) surface, which was tested due to its transparent properties in thin layers that could be applied to touchscreen surfaces. Whilst our study was ongoing, others reported different strategies that generated transparent surface coatings which exhibited significant anti-SARS-CoV-2 activity (33–35).

In addition to copper, we show that copper oxide (Cu_2_O-containing) test surfaces exhibited significant anti-SARS-CoV-2 activity, in agreement with other studies that have reported various copper oxide surfaces (CuO and/or Cu_2_O) to exhibit effective anti-SARS-CoV-2 activity (35–39). Importantly however, we demonstrate that the level of antiviral activity was strikingly dependent on the composition of carrier solution in which the virus inoculum was delivered to test surfaces. From our data, it can be concluded that when SARS-CoV-2 is delivered in DMEM-2%FBS (tissue culture media) a copper oxide surface (with Cu_2_O as the predominant oxidation phase) resulted in significantly better virus inactivation than the reference copper surface. However, the reverse is concluded when SARS-CoV-2 is delivered in AS (artificial saliva), as the reference copper surface exhibited superior antiviral activity over the Cu_2_O-containing surface. Further, SARS-CoV-2 delivered in PBS (phosphate buffered solution) resulted in the best virus inactivation whichever copper or Cu_2_O-containing surface was tested.

The purpose of evaluating antiviral activity of test surfaces in the laboratory is to identify surfaces that will perform efficiently in preventing fomite transmission when deployed on surfaces in public spaces. Therefore, although it is experimentally convenient to use a tissue culture derived virus inoculum (typically DEME or MEM with various FBS concentrations up to 10%) for evaluating the antiviral properties of test surfaces in the laboratory (5, 14–18, 35–39), we clearly demonstrate that antiviral performance of test surfaces is dependent upon the composition of the virus carrier solution. In real life, SARS-CoV-2 is expelled from an infected person via respiratory (saliva/sputum) droplets/aerosols, the composition of which is not accurately represented by tissue culture medium supplemented with FBS. In this study, we tested an artificial saliva carrier solution (40) formulated to mimic human saliva, which is a very dilute fluid composed of >97% water plus electrolytes, proteins/enzymes (41). Overall, our results suggest that future studies would ideally include virus delivered in physiologically relevant carrier solution, e.g., real human saliva/sputum samples when testing respiratory viruses, to recapitulate a real-life scenario to obtain a more realistic determination of test surface antiviral performance.

We hypothesized that composition of virus carrier solution could influence copper ion dissolution from copper/copper oxide surfaces, which would in turn effect surface antiviral performance if copper ion dissolution plays an important mechanistic role in antiviral activity. Indeed, we demonstrate that the different carrier solutions used in this study do influence the amount of copper ions released into solution from copper and Cu_2_O-containing surfaces, with the largest amount of copper ions released upon surface exposure to DMEM-2%FBS (tissue culture medium). In agreement with our observations, others have also reported that different liquids vary the level of ion release from copper and copper oxide surfaces and that the highest levels of release are into liquids containing amino acids, proteins or complex organic materials (42–44). Some studies have reported a positive correlation between the amount of copper ion released from copper/copper surfaces and antibacterial activity (45). However, we did not observe any correlation between SARS-CoV-2 inactivation and total amount of copper ions released in the presence of the different virus carrier solutions used in this study. Nevertheless, we proceeded to further investigate the role of copper ion dissolution, as our observation that surface thickness influenced level of antiviral activity also suggests that ion dissolution may play a mechanistic role in antiviral activity.

To do this, we performed a variation of the test surface inactivation assay, in which virus inactivation is de-coupled from the test copper/copper oxide surface to directly test if copper ions released into solution are capable of virus inactivation. Our results show that copper ions released into DMEM-2%FBS solution following copper or Cu_2_O-containing surface exposure, did not have the capacity to inactivate SARS-CoV-2. A reasonable interpretation of this observation could be that ion dissolution does not play a significant role in virus inactivation and instead direct surface contact killing is the major mechanism of action at play. Indeed, Hosseini *et al*., used a similar experimental approach to determine the role of copper ions released from a cupric oxide (CuO) film exposed to virus culture medium; material leached from their CuO coating did not inactivate SARS-CoV-2 and thus they rejected the hypothesis that dissolved material was the cause of inactivation and concluded that direct contact between SARS-CoV-2 and CuO is necessary to inhibit infection (36). Importantly, an alternative interpretation is required to explain our observations, because we provide direct evidence that copper ion dissolution is a mechanism by which SARS-CoV-2 can be inactivated, as material released into PBS solution following copper or Cu_2_O-containing surface exposure exhibited significant antiviral activity. The inactivation rate attributed to copper ion dissolution was ∼50% less than that observed when an equivalent virus inoculum was directly exposed to test surfaces, indicating that copper ion dissolution is not the only antiviral mechanism, and that direct contact killing may also play a role.

The question remains if copper ions released from our test surfaces are innately capable of virus inactivation, why doesn’t SARS-CoV-2 inactivation occur in DMEM-2%FBS solution released from our test surfaces? We propose that copper complexation with biomolecules (e.g., proteins, metabolites) in DMEM-2%FBS reduces the bioavailability of copper ions, therefore when SARS-CoV-2 is retrospectively added to released DMEM-2%FBS solution the copper ions are no longer available to interact with SARS-CoV-2 and thus virus inactivation does not occur. In support of this, Hedberg *et al*., demonstrated that copper ions released from Cu nanoparticles in biomolecule-containing media (e.g., DMEM, DMEM supplemented with FBS, or PBS supplemented with the amino acid histidine) does not exist as free Cu^2+^ ions in solution, but was instead completely complexed via strong bonds to biomolecules, conversely copper ions released from Cu nanoparticles in PBS formed labile Cu-complexes (42). Therefore, our interpretation of the data does not reject copper ion dissolution as an antiviral mechanism, on the contrary we provide direct evidence in support of copper ion dissolution as a mechanism that contributes to the antiviral activity of copper/copper oxide surfaces. Further, we suggest that complexation of dissolved copper ions with biomolecules present in the virus carrier solution can influence surface antiviral performance. We envision that competition between biomolecules in the carrier solution and the surface of SARS-CoV-2 for copper ion complexation could explain why our copper surfaces perform better when virus inoculum is delivered in PBS (which forms liable weak copper complexes) compared to DMEM-2%FBS (which forms strong chelating complexes) and further would explain why we did not observe a clear positive correlation between level of copper ion release into solution and antiviral activity. In support, whilst our manuscript was being prepared Glover *et al*., reported that coronavirus (OC43) inactivation on copper surfaces is significantly faster when virus was delivered in artificial perspiration solution compared to assay medium (DMEM) (44). Like our data, the rate of virus inactivation did not correlate with total amount of copper ions released into solution, instead they also suggest that chelated copper cations are not available for virus inactivation and that the organic constituents of DMEM act as chelators. Also, Sharan *et al*., who studied inactivation of *E.coli* suspensions in copper water storage vessels concluded that addition of amino acids, proteins or complex organic mixtures caused a dramatic decrease in *E.coli* inactivation, likely as a consequence of complex formation between leached copper and the organic constituents (43).

Further investigation is required to understand the results we obtained when virus was delivered in AS (which contains biomolecule mucin) (Fig.5), but our observations could suggest that different species of copper ions released from different copper surfaces could affect the degree of copper ion complexation and may also be dependent upon the type and level of chelating biomolecules present. For example, the presence of both ∼ 2 mM urea and the thiocynate ions in the AS will be forming various mixed hexadenate complexes with the copper ions in the aqueous solution.

It is also pertinent to stress that composition of virus carrier solution is just one important parameter, alongside multiple other variables that should be considered when assessing surface antiviral performance. Overall, these results further reiterate our conclusion that laboratory testing of surface antiviral performance should include virus delivered in a physiologically relevant carrier solution to replicate a real-life scenario and accurately assess the antiviral performance of test surfaces.

## Materials and Methods

### Generation of metal and metal oxide test surface coupons

All test surface coupons used are summarized in Tables S1 and S2. Thin film growth by electron beam evaporation was used to generated copper, bismuth and silver films. The films have been grown with various thicknesses using an e-beam evaporator in a vacuum of 10^-6^ mbar. Films were deposited with growth rates of approximately 10 nm/min with the substrate held at room temperature. Growth rates were calibrated using a quartz crystal microbalance. We employed silicon substrates (Inseto) with 5 nm thick nickel-chromium alloy (Ni-Cr) coating as an adhesion layer before growing the metals on top. The samples were cut into 4x4 mm^2^ pieces after growth to provide coupons for SARS-CoV-2 inactivation assays.

Thin film growth by molecular beam epitaxy was used to deposit palladium, nickel and transition metal oxide films. The films were grown using a reactive oxide molecular beam epitaxy system (MBE) (DCA Instruments Oy., Finland, dual R450), using thermal effusion cells, as well as an e-beam evaporator to evaporate the elemental metals. Metal films were grown in ultra-high vacuum at a pressure of ∼ 1 x 10^-9^ mbar, and oxide compounds in either molecular oxygen or 10 % ozone gas environment. Growth rates were calibrated using a quartz crystal microbalance prior to growth. Film thicknesses are controlled through the growth time. The typical pressure during growth varied from 2 x 10^-7^ to 2 x 10^-5^ mbar, depending on the materials. Samples were stored in a vacuum desiccator before use to avoid degradation due to exposure to air. Glass substrates (Nano Quartz Wafer GmbH, Germany) with a size of 4x4 mm^2^ were used for fabricating most of the surfaces, while aluminium oxide (Al_2_O_3_) (0001) substrates (MaTeck GmbH, Germany, of the same size) were used for copper chromate (CuCrO_2_). We have also used single crystalline substrates (LaAlO_3_)0.3(Sr_2_TaAlO_6_) (LSAT) (001), 4x4 mm^2^, from CrysTec GmbH, Germany) for identifying the crystalline phases using X-ray diffraction (XRD).

Thin film growth by magnetron sputtering was used to generate indium tin oxide (ITO) films, which were obtained from RF sputtering (Nexdep 030 DC/RF magnetron sputtering system, Angstrom Engineering Ltd., Canada) at 200 °C on glass substrates with a size of 4x4mm^2^. The total pressure of the argon environment was kept at ∼ 3 mTorr during the growth, and the films were annealed for 30 min after the sputtering.

For reference purposes we have included bulk foils of copper and stainless steel in the deactivation experiments. Copper foils of various thicknesses (Cu purity 99.9 %) and stainless steel (AISI 304) plates were obtained from GoodFellow Ltd., UK.

### Structural and morphological characterization of thin films

Film thicknesses were confirmed using a profilometer. X-ray diffraction (CuKα, 50 kV, Bruker Corp., USA, Discover D8) was used for phase identification of the materials and for obtaining crystallographic information such as grain size and orientation of the films (Fig. S1). To examine the elemental distribution along the thickness direction (Fig. S2), cross-sectional energy dispersive X-ray spectroscopy (EDX) (Thermo Fisher Scientific Inc., USA (formerly FEI) Titan Themis), performed in a transmission electron microscope (TEM), was utilised. The microscope was operated at 200 kV.

### Propagation of SARS-CoV-2 stocks

SARS-CoV-2 strain hCOV-19/England/2/2020 (kind gift of Dr Marian Killip, Public Health England, UK) was used within a class II Microbiology safety cabinet (MSC) inside a Biosafety Level 3 (BSL3) biocontainment facility. SARS-CoV-2 high titre stocks were propagated in Vero E6 cells (African green monkey kidney epithelial cell, ECACC, 85020206) as previously described (46). Briefly, Vero E6 cells were infected at a multiplicity of infection (MOI) of 0.01 and cells cultured in Dulbecco’s Modified Eagle’s Medium supplemented in 2% v/v FBS (DMEM-2%FBS) for 72 hours post-infection. Virus containing supernatant was collected and clarified by centrifugation for 15 mins at 3,200 x g at 4°C. The clarified stock was either directly aliquoted, flash frozen in liquid nitrogen and stored at -80°C or further concentrated using a polyethylene glycol (PEG) virus precipitation kit (Abcam) according to the manufacturer’s instructions. Briefly, 5 x PEG solution was mixed with clarified supernatant (1:4 ratio), incubated at 4°C overnight and then centrifuged at 3,200 x g for 30 minutes at 4°C. The resultant virus containing pellet was resuspended in DMEM-2%FBS using 1/100 volume of the starting virus supernatant and the resultant PEG-stock was then aliquoted, flash frozen in liquid nitrogen and stored at -80°C.

### Quantitation of viable SARS-CoV-2

Plaque assay was used, as previously described (46), to determine the titre of SARS-CoV-2 stocks as PFU/mL and to determine the % survival of SARS-CoV-2 following exposure to test antiviral surfaces. All plaques assays were performed in triplicate, plaques manually counted, followed by mean and SD determination. Plaque assay limit of detection (LOD) was determined via a 9-point 1:2 serial dilution of SARS-CoV-2 PEG-stock (that prior to the dilution series was diluted 1:1000 in DMEM-2%FBS to 5.8 x 10^5^ PFU/mL) to achieve theoretical zero. The SARS-CoV-2 plaque assay was performed, and plaque count plotted against dilution to generate a calibration curve. LOD was calculated with the following equation: LOD = 3 x (σ/S), with σ SD and S = slope of calibration curve R^2^= 0.9 (47).

### Test surface SARS-CoV-2 inactivation assay

Test surfaces (4x4 mm^2^ coupons) were disinfected in 70% v/v ethanol and allowed to air dry in a class II MSC for 15 minutes before transfer into 96-well plates using inverted forceps. A 7 μL droplet of SARS-CoV-2 virus inoculum containing ∼4000 PFU (derived from PEG-stock (5.8 x 10^8^ PFU/mL) diluted 1:1000 in DMEM-2%FBS) was pipetted onto the centre of each test coupon and incubated for the indicated times at room temperature. No coupon controls were conducted in parallel, were the equivalent 7 μL of virus inoculum was pipetted into sterile 1.5 mL tubes and incubated for the same length of time as corresponding inoculated test coupons. Recovery of virus from test surfaces was performed by adding 250 µL DMEM-2%FBS and gently pipetting up and down 25 times. The no coupon control was similarly processed. Recovered virus was transferred into individual wells of a 96 well plate and a 10-fold serial dilution in DMEM-2%FBS prepared to facilitate quantitation of % SARS-CoV-2 survival via plaque assay. To test the effect of various virus carrier solutions the SARS-CoV-2 PEG-stock (titre = 5.8 x 10^8^ PFU/mL) was utilized. In parallel, the PEG-stock stock was diluted 1:1000 in 3 different carrier virus solution (i) DMEM-2%FBS (ii) PBS or (iii) artificial saliva solution (AS; 0.18 mM MgCl_2_.7H_2_O, 1 mM CaCl_2_.H_2_O, 5 mM NaHCO_3_, 1.5 mM KH_2_PO_4_, 2.4 mM K_2_HPO_4_, 2 mM NH_4_Cl, 1.9 mM KSCN, 2 mM (NH_2_)_2_CO, 15 mM NaCl, 14 mM KCl, 0.3% w/v bovine salivary gland mucin (40). A droplet of 7 μL of virus in each carrier solution was verified by plaque assay to contain ∼4000 PFU and therefore the test surface virus inactivation assay was conducted as described above. The SARS-CoV-2 survival value was calculated as a percentage of the no coupon control sample and plotted as a bar chart as mean with SD and statistical significance assessed using two-way ANOVA with Tukey’s multiple comparison using Prism 9.5 GraphPad software. Significance is reported by P value *, p < 0.01, ***, p < 0.001, ****, p < 0.0001.

### De-coupled ion dissolution SARS-CoV-2 inactivation assay

To investigate the role of ion dissolution from test surfaces in SARS-CoV-2 inactivation, we performed a variation of the test surface inactivation assay described above, in which virus inactivation is decoupled from the test surface. Test surfaces were disinfected and dried as described above and 7 μL of each carrier solution (DMEM-2%FBS, PBS and AS) without virus, were pipetted onto the centre of each test coupon and incubated for 0 and 30 minutes at room temperature. No coupon controls were conducted in parallel, were the equivalent 7 μL of each carrier solution was pipetted into sterile 1.5 mL tubes and incubated for the same length of time as corresponding test coupons. After the indicated incubation times, the carrier solution was recovered from the test surface, transferred to 1.5mL tube and spiked with 2 μL of clarified SARS-CoV-2 stock (6 x 10^6^ PFU/mL) stock which had been diluted 1:3 in DMEM-2%FBS such that 2 μL contains ∼4000 PFU. SARS-CoV-2 incubation in each carrier solution pre-exposed to the test surfaces was performed for a further 0 or 30 minutes, followed by transfer into 96-well plates to facilitate quantitation of % SARS-CoV- 2 survival by plaque assay as described above.

### Inductively Coupled Plasma - Optical Emission Spectroscopy (ICP-OES)

ICP-OES was used to determine Cu ion concentration dissolved into carrier solution (DMEM-2%FBS, PBS or AS) upon exposure to various copper test surfaces. Prior to analysis test surfaces were disinfected and dried as described above and 7μL of each carrier solution added for 30 min at room temperature and then recovered in a further 250 μL of carrier solution, which was diluted 10X with 5% nitric acid. Alongside test samples, carrier solution controls not exposed to test surfaces, cupric acetate (∼90 ppm Cu ions) positive control and calibration standards (0, 0.005, 0.02 and 0.1 ppm Cu ions) were analysed. All analysis was conducted by The University of Edinburgh ICP analysis facility using a Vista-PRO Simultaneous ICP-OES (Varian/Agilent). LOD was calculated with the following equation: LOD = 3 x (σ/S), with σ = SD and S = slope of calibration curve R^2^= 0.9.

## Acknowledgments

This work was funded by UKRI-NIHR (MRC MR/V028464/1) COVID-19 Rapid Response Initiative. This grant was awarded to PW, CSA, TKS, PDCK, ADF. The funders have no role in the study, design, data collection and interpretation, or the decision to submit the work for publication. For the purpose of open access, the author(s) has applied a Creative Commons Attribution (CC BY) licence to any Accepted Manuscript version arising.

Author contributions: PW, CSA TKS, PDCK and ADF conceived the study. JH, YN, MF, MA, ADF, CSA executed the experiments. All authors analysed and interpreted the experimental data. CSA, PW, TKS, PDCK, JH, YN wrote the manuscript.

Underpinning data will made available at reference 48.

## Supplementary Material

**Table S1.** Summary of surfaces tested in this study as well as substrate materials, growth methods and profiles, and thickness.

| Tested Surfaces | Substrate | Growth method | Growth profile | Thickness (nm) |
| --- | --- | --- | --- | --- |
| Evaporated Copper (EC) | NiCr/Si | E-beam | Surface was deposited with e-beam in vacuum at room temperature after adhesive layer (Ni-Cr) deposition on Si. | 0 – 500 |
| Silver (Ag) | NiCr/Si | E-beam |  | 250 |
| Bismuth (Bi) | NiCr/Si | E-beam |  | 270 |
| Nickel (Ni) | Glass | MBE | Elemental metal sources were thermally evaporated in vacuum at room temperature directly on glass. | 30 |
| Palladium (Pd) | Glass | MBE |  | 20 |
| Copper Chromate ($\text{CuCrO}_2$) | $\text{Al}_2\text{O}_3$ | MBE | Cu and Cr was evaporated alternatively. $\text{O}_2$ pressure and substrate temperature was kept at $5 \times 10^{-6}$ mbar and at $800^\circ\text{C}$, respectively. | 25 |
| Indium Tin Oxide (ITO) | Glass | RF sputtering | ITO was sputtered at $200^\circ\text{C}$ with the total pressure of 3 mTorr, followed by post annealing for 30 min. | 10 |
| Titanium Oxide ($\text{TiO}_2$) | Glass | MBE | Ti was evaporated from effusion cell while $\text{O}_2$ pressure and substrate temperature were kept at $5 \times 10^{-6}$ mbar and at $700^\circ\text{C}$, respectively. | 16 |
| Annealed Evaporated Copper ($\text{CuO}/\text{Cu}_2\text{O}$) mixture | NiCr/Si | Annealing in air | Evaporated copper films were rinsed with acetone and 2-propanol for 5 min each, and dried at $200^\circ\text{C}$ for 5 min, then placed on hot surface at $350^\circ\text{C}$ in air and left for 1 hour. Cooled in air down to $200^\circ\text{C}$ then removed from the hot surface. Films were treated with water when required. | 100 |
|  |  |  |  | 500 |
| Copper Oxide ($\text{Cu}_2\text{O}$) | Glass LSAT | MBE | Cu was evaporated at $650^\circ\text{C}$ in 10 % $\text{O}_3$ environment where total pressure was kept at approx. $2 \times 10^{-5}$ mbar. | 10 |
|  |  |  |  | 30 |

**Table S2.** Summary of cutting, processing and storage of the surfaces tested in this study.

| Tested Surfaces | Cutting | Pre-processing | Storing after deposition |
| --- | --- | --- | --- |
| Evaporated copper (EC) | Manual cut with diamond pen after deposition | Rinsed with acetone then 2-propanol in ultra sonicator | Air |
| Silver (Ag) | (as above) | (as above) | (as above) |
| Bismuth (Bi) | (as above) | (as above) | (as above) |
| Nickel (Ni) | Precut substrates were used | (as above) | (as above) |
| Palladium (Pd) | (as above) | (as above) | (as above) |
| Copper Chromate (CuCrO<sub>2</sub>) | (as above) | (as above) | N<sub>2</sub> desiccator |
| Indium Tin Oxide (ITO) | Manual cut with diamond pen after deposition | (as above) | Air |
| Titanium Oxide (TiO<sub>2</sub>) | Precut substrates were used | (as above) | (as above) |
| Annealed Evaporated Copper (CuO/Cu<sub>2</sub>O) mixture | Cut EC coupons were used | Rinsed with water when needed | Vacuum storage |
| Copper Oxide (Cu<sub>2</sub>O) | Precut substrates were used | Rinsed with acetone then 2-propanol in ultra sonicator | (as above) |
| Copper foil | Cut with metal cutter | Polished one side, followed by rinsing with acetone, 2-propanol in ultra sonicator | Air |
| Stainless steel | (as above) | (as above) | (as above) |

**FIG S1.**
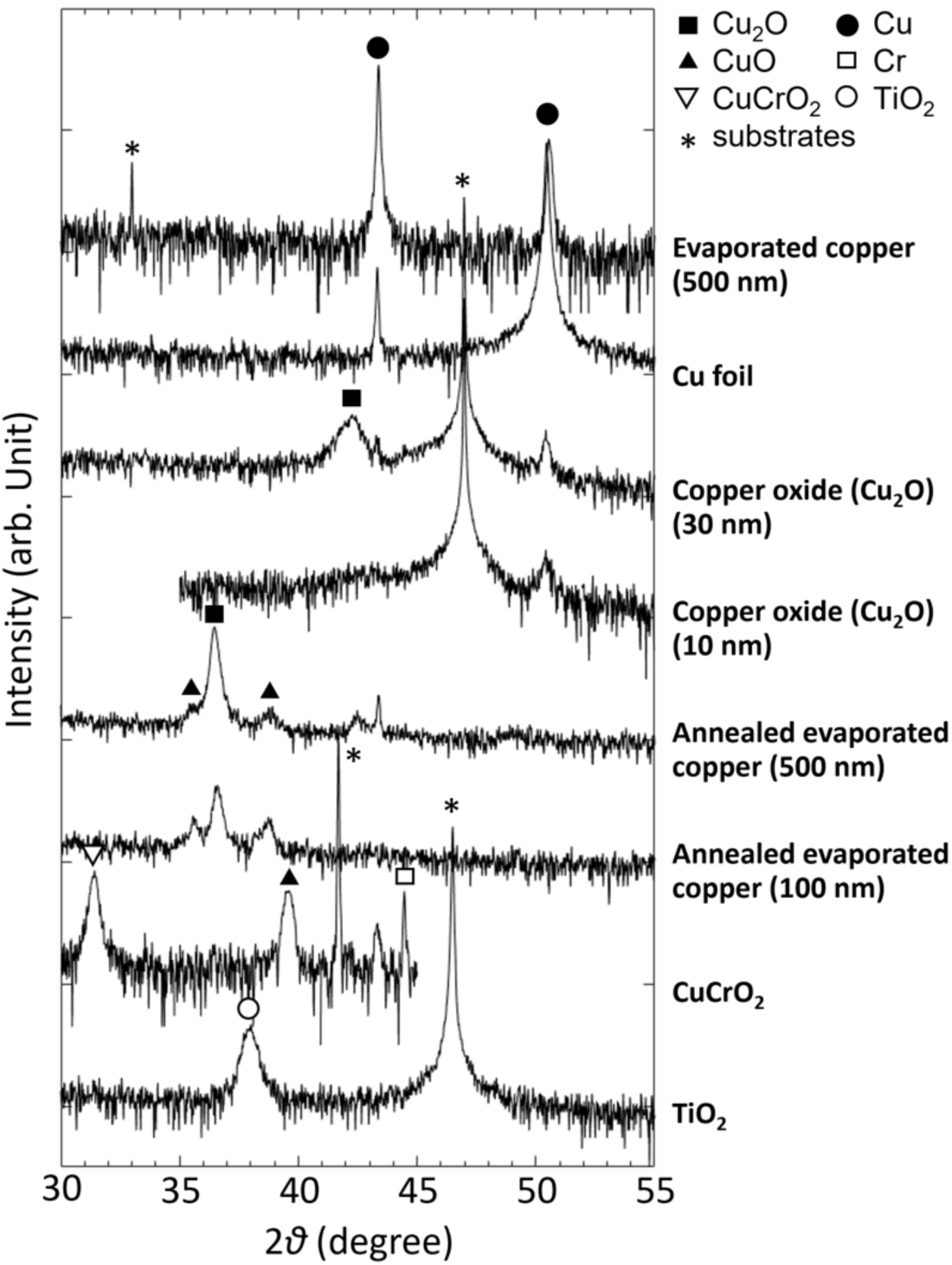
XRD patterns of surfaces tested in this study. Si (1 0 0) with adhesive layer of Ni-Cr alloy was used as a substrate material for films of evaporated copper and annealed copper, while Al_2_O_3_ (0 0 0 1), (LaAlO_3_)_0.3_(Sr_2_TaAlO_6_)_0.7_ (LSAT) (0 0 1), and SrTiO_3_ (0 0 1) were used for stabilising CuCrO_2_ and binary oxides, respectively. Diffraction peaks from substrate materials are all shown with asterisks (*). Note that the samples of copper oxide and titanium oxide used for virological tests were grown on glass substrates.

**FIG S2.**
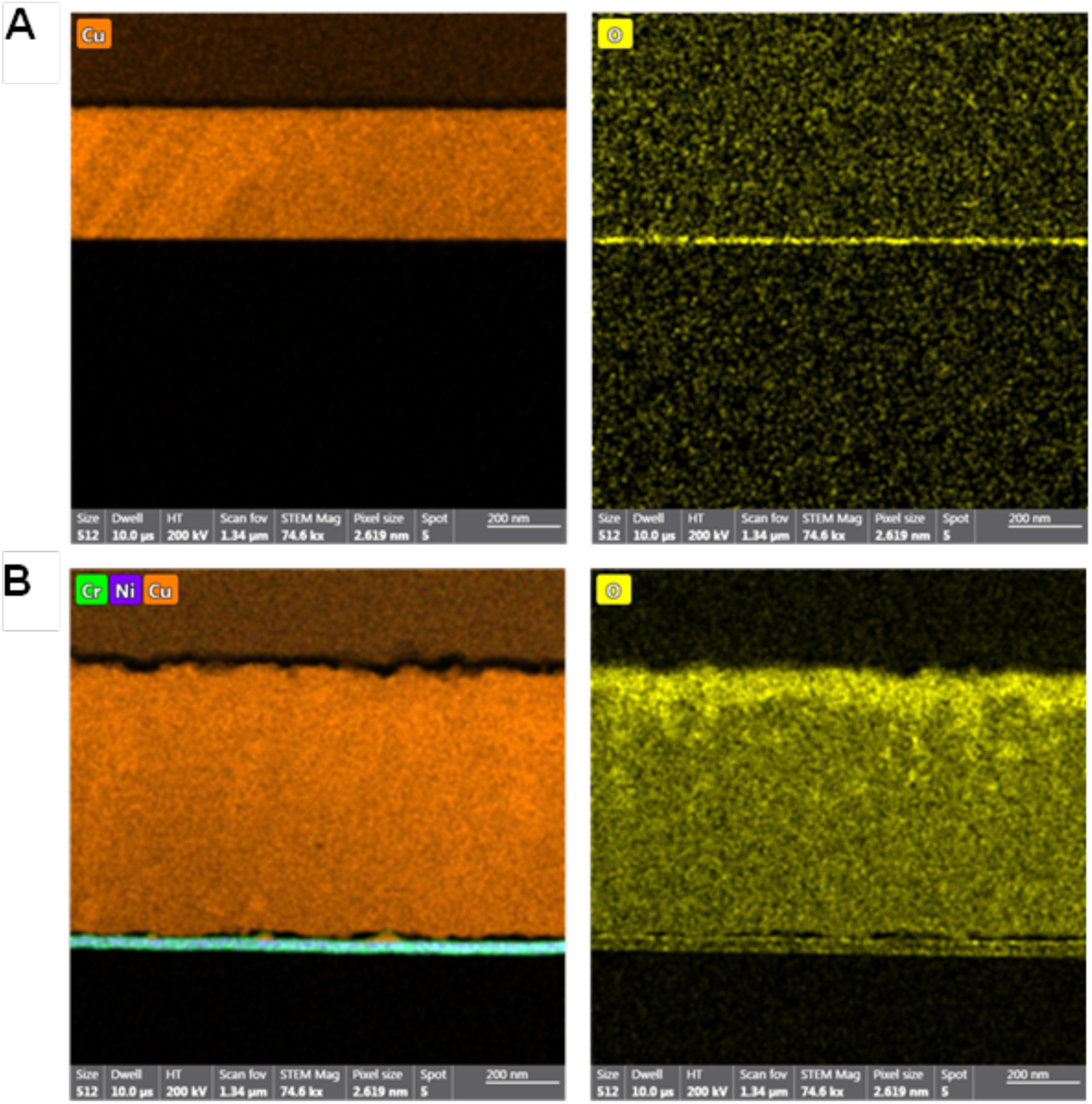
Cross-sectional energy dispersive X-ray analysis images on (A) evaporated copper film on crystalline Si, and (B) annealed evaporated copper. Smooth surface with sharp interface is apparent in images from evaporated copper while the evaporated copper film show rougher surface. Notably, the distribution of oxygen atoms in the annealed copper film (bottom right) is not uniform and higher density of oxygen near the film surface can be seen.

**FIG S3.**
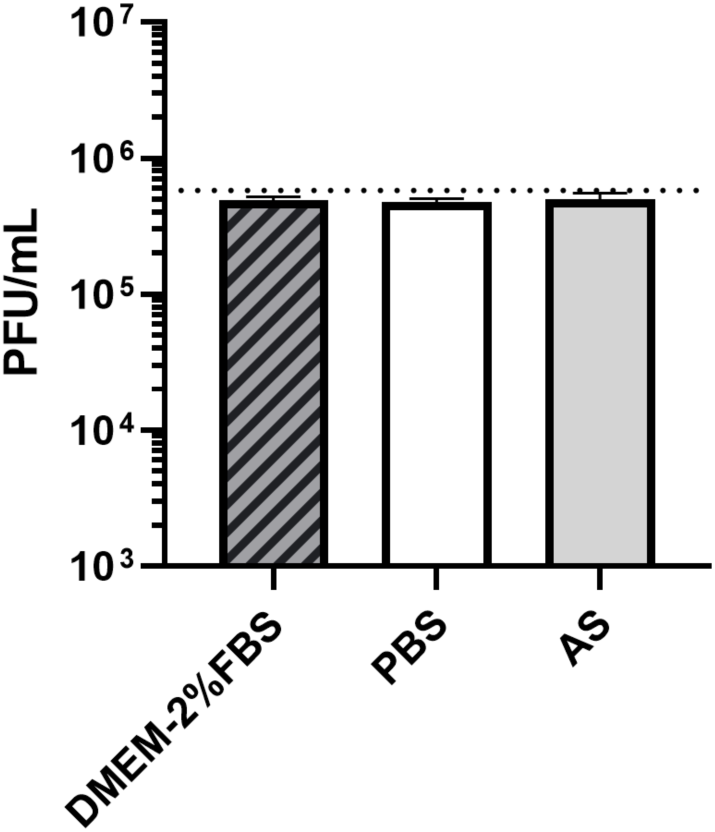
Titre of SARS-CoV-2 in Carrier solutions. Following resuspension of SARS-CoV-2 in each carrier solution, the virus was confirmed as remaining viable. SARS-CoV-2 viral titre was determined, and data is presented as (PFU/mL) in each carrier solution, DMEM-2%FBS, PBS and AS. Data shown represents mean values (n = 3 replicates and error bar = SD) and is representative of 3 independent experiments.

## References

1. Birkett M, Dover L, Cherian Lukose C, Wasy Zia A, Tambuwala MM, Serrano-Aroca A. 2022. Recent Advances in Metal-Based Antimicrobial Coatings for High-Touch Surfaces. Int J Mol Sci 23.

2. Boone SA, Gerba CP. 2007. Significance of fomites in the spread of respiratory and enteric viral disease. Appl Environ Microbiol 73:1687–96.

3. Leung NHL. 2021. Transmissibility and transmission of respiratory viruses. Nat Rev Microbiol 19:528–545.

4. Geng Y, Wang Y. 2023. Stability and transmissibility of SARS-CoV-2 in the environment. J Med Virol 95:e28103.

5. van Doremalen N, Bushmaker T, Morris DH, Holbrook MG, Gamble A, Williamson BN, Tamin A, Harcourt JL, Thornburg NJ, Gerber SI, Lloyd-Smith JO, de Wit E, Munster VJ. 2020. Aerosol and Surface Stability of SARS-CoV-2 as Compared with SARS-CoV-1. N Engl J Med 382:1564–1567.

6. Pastorino B, Touret F, Gilles M, de Lamballerie X, Charrel RN. 2020. Prolonged Infectivity of SARS-CoV-2 in Fomites. Emerg Infect Dis 26.

7. Zhou J, Otter JA, Price JR, Cimpeanu C, Meno Garcia D, Kinross J, Boshier PR, Mason S, Bolt F, Holmes AH, Barclay WS. 2021. Investigating Severe Acute Respiratory Syndrome Coronavirus 2 (SARS-CoV-2) Surface and Air Contamination in an Acute Healthcare Setting During the Peak of the Coronavirus Disease 2019 (COVID-19) Pandemic in London. Clin Infect Dis 73:e1870–e1877.

8. Colaneri M, Seminari E, Novati S, Asperges E, Biscarini S, Piralla A, Percivalle E, Cassaniti I, Baldanti F, Bruno R, Mondelli MU, Force CISMPT. 2020. Severe acute respiratory syndrome coronavirus 2 RNA contamination of inanimate surfaces and virus viability in a health care emergency unit. Clin Microbiol Infect 26:1094 e1–1094 e5.

9. Santarpia JL, Rivera DN, Herrera VL, Morwitzer MJ, Creager HM, Santarpia GW, Crown KK, Brett-Major DM, Schnaubelt ER, Broadhurst MJ, Lawler JV, Reid SP, Lowe JJ. 2020. Aerosol and surface contamination of SARS-CoV-2 observed in quarantine and isolation care. Sci Rep 10:12732.

10. Ahn JY, An S, Sohn Y, Cho Y, Hyun JH, Baek YJ, Kim MH, Jeong SJ, Kim JH, Ku NS, Yeom JS, Smith DM, Lee H, Yong D, Lee YJ, Kim JW, Kim HR, Hwang J, Choi JY. 2020. Environmental contamination in the isolation rooms of COVID-19 patients with severe pneumonia requiring mechanical ventilation or high-flow oxygen therapy. J Hosp Infect 106:570–576.

11. Grass G, Rensing C, Solioz M. 2011. Metallic copper as an antimicrobial surface. Appl Environ Microbiol 77:1541–7.

12. Ramos-Zuniga J, Bruna N, Perez-Donoso JM. 2023. Toxicity Mechanisms of Copper Nanoparticles and Copper Surfaces on Bacterial Cells and Viruses. Int J Mol Sci 24.

13. Salah I, Parkin IP, Allan E. 2021. Copper as an antimicrobial agent: recent advances. RSC Adv 11:18179–18186.

14. Liu LT, Chin AWH, Yu P, Poon LLM, Huang MX. 2022. Anti-pathogen stainless steel combating COVID-19. Chem Eng J 433:133783.

15. Mosselhy DA, Kareinen L, Kivisto I, Aaltonen K, Virtanen J, Ge Y, Sironen T. 2021. Copper-Silver Nanohybrids: SARS-CoV-2 Inhibitory Surfaces. Nanomaterials (Basel) 11.

16. Meister TL, Fortmann J, Breisch M, Sengstock C, Steinmann E, Koller M, Pfaender S, Ludwig A. 2022. Nanoscale copper and silver thin film systems display differences in antiviral and antibacterial properties. Sci Rep 12:7193.

17. Mantlo EK, Paessler S, Seregin A, Mitchell A. 2021. Luminore CopperTouch Surface Coating Effectively Inactivates SARS-CoV-2, Ebola Virus, and Marburg Virus In Vitro. Antimicrob Agents Chemother 65:e0139020.

18. Hutasoit N, Kennedy B, Hamilton S, Luttick A, Rahman Rashid RA, Palanisamy S. 2020. Sars-CoV-2 (COVID-19) inactivation capability of copper-coated touch surface fabricated by cold-spray technology. Manuf Lett 25:93–97.

19. Terzioglu E, Arslan M, Balaban BG, Cakar ZP. 2022. Microbial silver resistance mechanisms: recent developments. World J Microbiol Biotechnol 38:158.

20. Manakhov AM, Permyakova ES, Sitnikova NA, Tsygankova AR, Alekseev AY, Solomatina MV, Baidyshev VS, Popov ZI, Blahova L, Elias M, Zajickova L, Kovalskii AM, Sheveyko AN, Kiryukhantsev-Korneev PV, Shtansky DV, Necas D, Solovieva AO. 2022. Biodegradable Nanohybrid Materials as Candidates for Self-Sanitizing Filters Aimed at Protection from SARS-CoV-2 in Public Areas. Molecules 27.

21. Delumeau LV, Asgarimoghaddam H, Alkie T, Jones AJB, Lum S, Mistry K, Aucoin MG, DeWitte-Orr S, Musselman KP. 2021. Effectiveness of antiviral metal and metal oxide thin-film coatings against human coronavirus 229E. APL Mater 9:111114.

22. Jeremiah SS, Miyakawa K, Morita T, Yamaoka Y, Ryo A. 2020. Potent antiviral effect of silver nanoparticles on SARS-CoV-2. Biochem Biophys Res Commun 533:195–200.

23. Galdiero S, Falanga A, Vitiello M, Cantisani M, Marra V, Galdiero M. 2011. Silver nanoparticles as potential antiviral agents. Molecules 16:8894–918.

24. Kumar A, Nath K, Parekh Y, Enayathullah MG, Bokara KK, Sinhamahapatra A. 2021. Antimicrobial silver nanoparticle-photodeposited fabrics for SARS-CoV-2 destruction. Colloid Interface Sci Commun 45:100542.

25. He Q, Lu J, Liu N, Lu W, Li Y, Shang C, Li X, Hu L, Jiang G. 2022. Antiviral Properties of Silver Nanoparticles against SARS-CoV-2: Effects of Surface Coating and Particle Size. Nanomaterials (Basel) 12.

26. Baselga M, Uranga-Murillo I, de Miguel D, Arias M, Sebastian V, Pardo J, Arruebo M. 2022. Silver Nanoparticles-Polyethyleneimine-Based Coatings with Antiviral Activity against SARS-CoV-2: A New Method to Functionalize Filtration Media. Materials (Basel) 15.

27. Bono N, Ponti F, Punta C, Candiani G. 2021. Effect of UV Irradiation and TiO2-Photocatalysis on Airborne Bacteria and Viruses: An Overview. Materials (Basel) 14.

28. Micochova P, Chadha A, Hesseloj T, Fraternali F, Ramsden JJ, Gupta RK. 2021. Rapid inactivation of SARS-CoV-2 by titanium dioxide surface coating. Wellcome Open Res 6:56.

29. Nakano R, Yamaguchi A, Sunada K, Nagai T, Nakano A, Suzuki Y, Yano H, Ishiguro H, Miyauchi M. 2022. Inactivation of various variant types of SARS-CoV-2 by indoor-light-sensitive TiO2-based photocatalyst. Sci Rep 12:5804.

30. Matsuura R, Lo CW, Wada S, Somei J, Ochiai H, Murakami T, Saito N, Ogawa T, Shinjo A, Benno Y, Nakagawa M, Takei M, Aida Y. 2021. SARS-CoV-2 Disinfection of Air and Surface Contamination by TiO2 Photocatalyst-Mediated Damage to Viral Morphology, RNA, and Protein. Viruses 13.

31. Han R, Coey JD, O’Rourke C, Bamford CGG, Mills A. 2022. Flexible, disposable photocatalytic plastic films for the destruction of viruses. J Photochem Photobiol B 235:112551.

32. Lu Y, Guan S, Hao L, Yoshida H, Nakada S, Takizawa T, Itoi T. 2022. Inactivation of SARS-CoV-2 and photocatalytic degradation by TiO2 photocatalyst coatings. Sci Rep 12:16038.

33. Hosseini M, Chin AWH, Williams MD, Behzadinasab S, Falkinham JO, 3rd, Poon LLM, Ducker WA. 2022. Transparent Anti-SARS-CoV-2 and Antibacterial Silver Oxide Coatings. ACS Appl Mater Interfaces 14:8718–8727.

34. Gentili V, Pazzi D, Rizzo S, Schiuma G, Marchini E, Papadia S, Sartorel A, Di Luca D, Caccuri F, Bignozzi CA, Rizzo R. 2021. Transparent Polymeric Formulations Effective against SARS-CoV-2 Infection. ACS Appl Mater Interfaces 13:54648–54655.

35. Behzadinasab S, Williams MD, Hosseini M, Poon LLM, Chin AWH, Falkinham JO, 3rd, Ducker WA. 2021. Transparent and Sprayable Surface Coatings that Kill Drug-Resistant Bacteria Within Minutes and Inactivate SARS-CoV-2 Virus. ACS Appl Mater Interfaces 13:54706–54714.

36. Hosseini M, Chin AWH, Behzadinasab S, Poon LLM, Ducker WA. 2021. Cupric Oxide Coating That Rapidly Reduces Infection by SARS-CoV-2 via Solids. ACS Appl Mater Interfaces 13:5919–5928.

37. Behzadinasab S, Chin A, Hosseini M, Poon L, Ducker WA. 2020. A Surface Coating that Rapidly Inactivates SARS-CoV-2. ACS Appl Mater Interfaces 12:34723–34727.

38. Merkl P, Long S, McInerney GM, Sotiriou GA. 2021. Antiviral Activity of Silver, Copper Oxide and Zinc Oxide Nanoparticle Coatings against SARS-CoV-2. Nanomaterials (Basel) 11.

39. Purniawan A, Lusida MI, Pujiyanto RW, Nastri AM, Permanasari AA, Harsono AAH, Oktavia NH, Wicaksono ST, Dewantari JR, Prasetya RR, Rahardjo K, Nishimura M, Mori Y, Shimizu K. 2022. Synthesis and assessment of copper-based nanoparticles as a surface coating agent for antiviral properties against SARS-CoV-2. Sci Rep 12:4835.

40. Woo MH, Hsu YM, Wu CY, Heimbuch B, Wander J. 2010. Method for contamination of filtering facepiece respirators by deposition of MS2 viral aerosols. J Aerosol Sci 41:944–952.

41. Diaz-Arnold AM, Marek CA. 2002. The impact of saliva on patient care: A literature review. J Prosthet Dent 88:337–43.

42. Hedberg J, Karlsson HL, Hedberg Y, Blomberg E, Odnevall Wallinder I. 2016. The importance of extracellular speciation and corrosion of copper nanoparticles on lung cell membrane integrity. Colloids Surf B Biointerfaces 141:291–300.

43. Sharan R, Chhibber S, Attri S, Reed RH. 2010. Inactivation and sub-lethal injury of Escherichia coli in a copper water storage vessel: effect of inorganic and organic constituents. Antonie Van Leeuwenhoek 98:103–15.

44. Glover CF, Miyake T, Wallemacq V, Harris JD, Emery J, Engel DA, McDonnell SJ, Scully JR. 2022. Interrogating the Effect of Assay Media on the Rate of Virus Inactivation of High-Touch Copper Surfaces: A Materials Science Approach. Advanced Materials Interfaces 9.

45. Hans M, Erbe A, Mathews S, Chen Y, Solioz M, Mucklich F. 2013. Role of copper oxides in contact killing of bacteria. Langmuir 29:16160–6.

46. Gruschow S, Adamson CS, White MF. 2021. Specificity and sensitivity of an RNA targeting type III CRISPR complex coupled with a NucC endonuclease effector. Nucleic Acids Res 49:13122–13134.

47. Sengul U. 2016. Comparing determination methods of detection and quantification limits for aflatoxin analysis in hazelnut. J Food Drug Anal 24:56–62.

48. 48. Hilton, J., et al. Dataset for "", St Andrews research repository, <DOI inserted on acceptence> (2023).

